# RhoV Promotes Proliferation, Migration, and MAPK Activation in Pancreatic Ductal Adenocarcinoma

**DOI:** 10.1101/2025.09.10.672939

**Authors:** Shang Wu, Jia Feng, Ting Wang, Kyler Hwa, Selina Kao, Ying Xiao, Dongfang Yang, Xinjian Li, Zebang Li, Ryan Liu, Yancheng Liu, Yikai Lu, Xiaoxu Zhou, Yifei Liu, Chiung-Kuei Huang, Shaolei Lu

## Abstract

**Background and Aims:** Pancreatic ductal adenocarcinoma (PDAC) is a highly lethal malignancy characterized by diagnosis at advanced stages, limited therapeutic options, and frequent resistance to therapies. Although oncogenic KRAS mutations are central drivers of PDAC, alternative pathways indisputably contribute to its tumorigenesis and progression. RhoV, a member of the Rho family of small GTPases, has been implicated in tumor development in other cancer types, such as breast cancer and lung adenocarcinoma; however, its role in PDAC remains unclear.

**Methods:** In this study, we investigated the expression and functional impact of RhoV on PDAC. Analysis of publicly available datasets and immunohistochemical profiling of 114 PDAC patient specimens were used to evaluate the expression of RhoV in PDAC and its prognostic impact. Overexpression and CRISPR-Cas9-mediated knockout of RhoV were established in three pancreatic cancer cell lines. Functional analyses, such as cell proliferation, migration, invasion, colony formation, spheroid growth, and mouse xenograft, were used to evaluate the role of RhoV in PDAC cells.

**Results:** RhoV overexpression was associated with reduced overall and recurrence-free survival in public datasets and our own patient cohort. Functional assays demonstrated that RhoV overexpression promoted PDAC cell proliferation, colony formation, and spheroid growth, whereas knockout of RhoV suppressed these changes. Moreover, RhoV enhanced PDAC cell migration and invasion *in vitro*, accompanied by downregulation of E-cadherin and upregulation of N-cadherin and vimentin, indicating induction of epithelial–mesenchymal transition. Mechanistically, RhoV overexpression activated key MAPK pathway components, including phosphorylation of ERK, JNK, and p38. *In vivo*, xenograft models confirmed that RhoV drives tumor growth and increases tumor burden.

**Conclusion:** These results establish RhoV as a novel oncogenic factor in PDAC progression and highlight its potential as a biomarker and therapeutic target, warranting further investigation into combinatorial targeting strategies to overcome KRAS inhibitor resistance.

## Introduction

Pancreatic cancer ranks among the top causes of cancer-related deaths in developed countries and stands as one of the deadliest malignant tumors globally [1]. The vast majority, about 95% of pancreatic cancers, originate from the exocrine part of the pancreas. Among these, the predominant type is pancreatic ductal adenocarcinoma (PDAC), which is an extremely aggressive and often fatal form of cancer [2]. Despite constituting only 3% of new cancer diagnoses in the United States, PDAC is projected to become the second leading cause of cancer-related deaths by 2030 [3]. The prognosis for pancreatic cancer is generally poor, with median survival times of approximately 10-12 months even with treatment [4]. This is largely due to the advanced stage at which the diagnosis is made, and only about 10–20% are candidates for surgical resection [5]. For patients with locally unresectable or metastatic adenocarcinoma, chemotherapy has been shown to improve the overall 1-year survival rate without significantly enhancing 5-year survival rate [6, 7]. The most frequently mutated gene in PDAC is the oncogene KRAS [8, 9]. Notably, about 90% of individuals with PDAC possess the G12 mutation in the KRAS codon [10]. Unfortunately, no direct inhibitors of KRAS or its mutant forms have been approved for clinical use in PDAC patients. This challenge is partly attributed to the complex nature of KRAS signaling, which can activate multiple alternative pathways [11]. When KRAS is abnormally activated, it can initiate signaling through various pathways, including the MAPK and mTOR pathways, as well as interact with other molecules like RALGDS, TIAM1, and RIN1[12]. This complexity complicates the efficacy of KRAS inhibitors, even if inhibitors of specific KRAS mutants have recently been developed. Finding new therapeutic targets outside of the KRAS pathway has become increasingly imperative.

All Ras-like proteins belong to a large superfamily that is divided into five main families: Ras, Rho, Rab, Arf, and Ran. Each family specifically regulates different aspects of cell metabolism, highlighting their diverse and crucial roles in cellular function [13]. Rho GTPases constitute a unique family within the Ras superfamily of small GTP-binding proteins. RhoV belongs to the Rho family and plays a crucial role in neural crest differentiation and cell development [14, 15]. Existing research has demonstrated RhoV mRNA dysregulation across a range of tumors, with elevated levels in 18 different tumor types, including breast carcinoma, lung adenocarcinoma, and pancreatic adenocarcinoma [16, 17]. For instance, RhoV is upregulated in non-small cell lung cancer and affects cancer progression [18] and contributes to tumor growth and EMT via JNK phosphorylation in adenocarcinoma [19]. Another study showed that knockdown of RhoV inhibits the activation of the downstream EGFR signaling pathway, including the phosphorylation of ERK and AKT [20]. In triple-negative breast cancer, RhoV also plays an important role in metastasis by binding EGFR to initiate the JNK pathway [21]. However, the role and function of RhoV in pancreatic adenocarcinoma are still insufficiently investigated.

In this study, we reported that RhoV expression was associated with poor prognosis and survival rate. We investigated the effects of RhoV on pancreatic adenocarcinoma in vivo and in vitro by overexpressing and knocking out the RhoV gene. The underlying mechanism showed that RhoV is involved in the MAPK pathway and promotes phosphorylation of key regulators in the pathway.

## Methods and Materials

### Patients and specimens

A total of 114 cases of PDAC from pancreaticoduodenectomy or distal pancreatectomy) were collected from the pathology archive of Rhode Island Hospital between 2010 and 2023. Ampullary and duodenal carcinomas were excluded. Clinicopathological parameters, such as survival data, were retrieved from the hospital cancer registry and/or medical chart review. The study was approved by the Internal Review Board of Brown University Health.

### Immunohistochemistry

Representative whole sections of each PDAC were selected to perform the immunohistochemical stains. Briefly, 4-μm paraffin sections were deparaffinized with xylene and rehydrated in graded alcohols and water. Heat-Induced Epitope Retrieval (HIER) was achieved by boiling the slides in citrate buffer (pH 6.0) for 30 minutes. Endogenous peroxidase activity was quenched using Dako Dual Endogenous Enzyme Block (Agilent). The sections were then incubated with a rabbit polyclonal anti-RhoV antibody (26620-1-AP, 1:400, Protentech, Rosemont, IL) overnight at 4°C. The immunoreactivity was detected using a 3,3’-diaminobenzidine (DAB) Substrate Chromogen System (Agilent). Immunohistochemical staining for RhoV was evaluated individually to give staining intensity and the proportion of positive tumor cells (by S.L.). At least mild staining intensity in at least 5% of the tumor cells was considered high expression.

### Human cancer cell lines

Human cell lines HKT-293T and pancreatic cancer cell lines PANC1, AsPc1, and HPAF2 were purchased from the American Type Culture Collection, ATCC. The cell lines HKT-293T, PANC1, and HPAF2 were grown in Dulbecco’s modified Eagle’s medium (DMEM) with 10% fetal bovine serum (FBS), 1% L-glutamine, 1% penicillin. AsPc1 cells were grown in Roswell Park Memorial Institute medium (RPMI 1640) with 10% fetal bovine serum, 1% L-glutamine, 1% penicillin. All of the cells were cultured in a humidified incubator at 37°C and 5% CO_2_.

### Virus production and infection

For overexpressed RhoV, RhoV lentiviral vector, psPAX2, pMD2G plasmids were used. For knockout RhoV, RhoV sgRNA CRISPR/Cas9 All-in-One lentivector, psPAX2, pMD2G plasmids were used. The plasmids were transfected into HEK293T cells with 8.5 μL polyplus transfection reagent for lentiviral generation. HEK293T packaging cells were plated into 10 cm plates in DMEM 24 hours before transfection. After transfection, the lentivirus supernatant was collected at 48 hours and 72 hours. The supernatant was filtered through a 0.22 μm PES filter. The lentiviral stock was diluted with cell culture medium (ratio 1:1 or 1:2) and transduced target cells in the presence of polybrene (8 μg/ml). 72 hours post-lentivirus infection, the cells were ready for selection and experiments. Successfully infected PANC1 cells were selected with puromycin (2-2.5 μg/ml). HPAF2 cells were selected with puromycin (0.5-1.5 μg/ml). AsPc1 cells were selected with puromycin (1.5 μg/ml). RhoV lentiviral vector (Cat. No.395310610295) and RhoV sgRNA CRISPR/Cas9 All-in-One lentivector (Cat. No.39531111) plasmids were purchased from Applied Biological Materials (Richmond, Canada).

### Quantitative reverse transcription polymerase chain reaction (RT-qPCR)

For RT-qPCR, RNA was extracted from cells by TRIzol (Invitrogen) and iScript™ cDNA Synthesis Kit (Bio-Rad) to transform 1 μg RNA into cDNA. cDNA was mixed with SYBR green, water, and primer. Samples were processed using QuantStudio5 Real-Time PCR System. Specific primers were used for RhoV (5-GGGAACGAAGTCGGTCAAA-3; 5-ACACCTTCTCTGTGCAAGTC-3; 5-GGTAGCAAAGGGAACGAAGT-3; 5-GCAAGTCCTGGTGGATGG-3) and 18S (5-GAGACTCTGGCATGCTAACTAG-3; 5-GGACATCTAAGGGCATCACAG-3), with GAPDH (5-GATGGCAACAATATCCACTTTACC-3; 5-GGTCGGAGTTCAACGGATTT-3) as reference control.

### Western blotting

RIPA buffer with 1% protease and phosphatase inhibitors was used to collect whole cell extracts. The cell lysate was dissolved in the 3×SDS-PAGE loading buffer and boiled for 10 minutes. The proteins were separated by 8% or 10% SDS-PAGE gel and transferred to PVDF membrane. The membrane was pre-treated with 5% milk or 4% BSA in TBST solution for 1 hour, followed by incubating with the primary antibodies at 4°C overnight: mouse polyclonal anti-GAPDH (ABclonal; Cat.No. AC033, 1:2000), rabbit polyclonal anti-RhoV (Osenses; OSR00321W, 1:1000), rabbit polyclonal anti-p38 (Cell Signaling Technology [CST]; #9212, 1:1000), rabbit polyclonal anti-Phospho-p38 (CST; #9211S, 1:1000), rabbit polyclonal anti-ERK (CST; #4695S, 1:1000), rabbit polyclonal anti-Phospho-ERK (CST; #4370S, 1:1000), rabbit polyclonal anti-SAPK/JNK (CST; #9252, 1:1000), rabbit polyclonal anti-Phospho-SAPK/JNK (CST; #9251, 1:1000), rabbit polyclonal anti-N-Cadherin (CST; #13116, 1:1000), rabbit polyclonal anti-E-Cadherin (CST; #3195, 1:1000), rabbit polyclonal anti-Vimentin (CST; #5741, 1:1000), rabbit polyclonal anti-α-tubulin (CST; #80762, 1:2000). After washing in TBST for 10 minutes, the membrane was incubated for 60 mins at room temperature with HRP-conjugated goat anti-mouse antibody (Vector Laboratories; 1:10000) or goat anti-rabbit antibody (Vector Laboratories; 1:10000), and individual protein band was visualized with chemiluminescence substrate (Thermo Fisher Scientific) at ChemiDoc Imaging System (Bio-Rad, Hercules, CA)

### Colony formation assay

500 cells were plated in the 6-well plate. The culture medium was replaced twice a week. After 2-3 weeks, the colonies were fixed with 10% formaldehyde for 20 mins, followed by 10% Giemsa dye to stain the colonies.

### Tumor spheroid formation assay

500 cells were plated in the 96-well round-bottom ultra-low attachment plate. On the seventh day, the tumor spheroids were observed under a 4× magnification microscope. ImageJ was utilized to measure the diameters of the tumor spheroids.

### MTT assay

Cells were seeded in a 96-well plate with 500 cells per well. MTT stock solution was diluted with culture medium at a ratio of 1:10 to prepare the working solution. At different time points, the MTT working solution was added to the wells and incubated for one hour. The resulting crystals were dissolved with DMSO, and absorbance was measured using a microplate reader at 490 nm.

### Scratch assay

Cells were seeded on a 6-well plate with 3×10^6^ in serum-free medium. After 24 hours, uniform scratches were made in the monolayers, followed by washing and adding fresh medium. The wound areas were monitored under the same microscope at zero, 24, and 48 hours.

### Transwell assay

24-well transwell plates were used for the invasion assay. Transwell plate membranes were pre-coated with diluted Matrigel (12.5%; Corning, USA), and cells (5×10^4^ cells in serum-free medium) were plated in the upper chamber while 10% serum-containing media was placed in the lower chamber. After 48h of incubation, cells that invaded the lower chamber were fixed with methanol for 30 minutes. The cells were stained with 0.05% crystal violet for 10 minutes, then washed three times with PBS, and positively stained cells were counted and observed under a 10× magnification microscope. ImageJ is used to quantify the region of invasion.

### Animal model

For the xenograft mouse model, female NOD/SCID mice (6–8 weeks old) purchased from Jackson Lab (Bar Harbor, ME) were maintained under specific-pathogen-free conditions. Nude mice were inoculated subcutaneously (s.c.) in the lower right flank with 1×10^7^ cells mixed with Matrigel matrix and serum-free medium (ratio of 1:1) in a total volume of 100 μL. All protocols were approved by the Institutional Animal Care and Use Committee (IACUC) at Tulane University, and all experiments were conducted in accordance with the guidelines of this IACUC.

### Statistical analysis

Statistical analysis was performed with GraphPad Prism 8.0 or 10.0 (GraphPad Software). Student t-test or Analysis of Variance (ANOVA) was used, and p-values less than 0.05 were considered significant.

## Results

### RhoV is associated with patients’ survival and prognosis in pancreatic adenocarcinoma

In the pan-cancer analysis, RhoV was significantly elevated in pancreatic adenocarcinoma, especially enriched in the basal-like type [17] [22]. Based on the Protein Atlas database (proteinatlas.org), the gene expression of RhoV is significantly associated with worse prognosis of pancreatic cancer patients (P<0.0001, Figure 1A, accessed in July 2025). Subsequently, we performed RhoV immunostaining on 114 pancreatic cancer samples from patients. RhoV showed a typical cytoplasmic staining pattern. No other cells or tissues exhibited any staining except nonspecific stains in the necrotic debris (Figure 1B). In the Kaplan-Meier chart analysis of the patient outcome, we found that high expression of RhoV (at least mildly expressed in at least 5% of tumor cells) was negatively correlated with both overall survival and recurrence-free survival (Figure 1C, P=0.0182 and Figure 1D, P=0.0198). Both external and internal data confirmed that RhoV was related to patients’ poor survival rates. Diverse expression of RhoV was also observed among human PDAC cell lines. We examined the mRNA and protein levels of RhoV in three human PDAC cell lines that we have, and found that HPAF2 had relatively higher levels of RhoV, while AsPc1 had lower levels, compared to PANC1 (Figure 1E and 1F).

**Figure 1.**
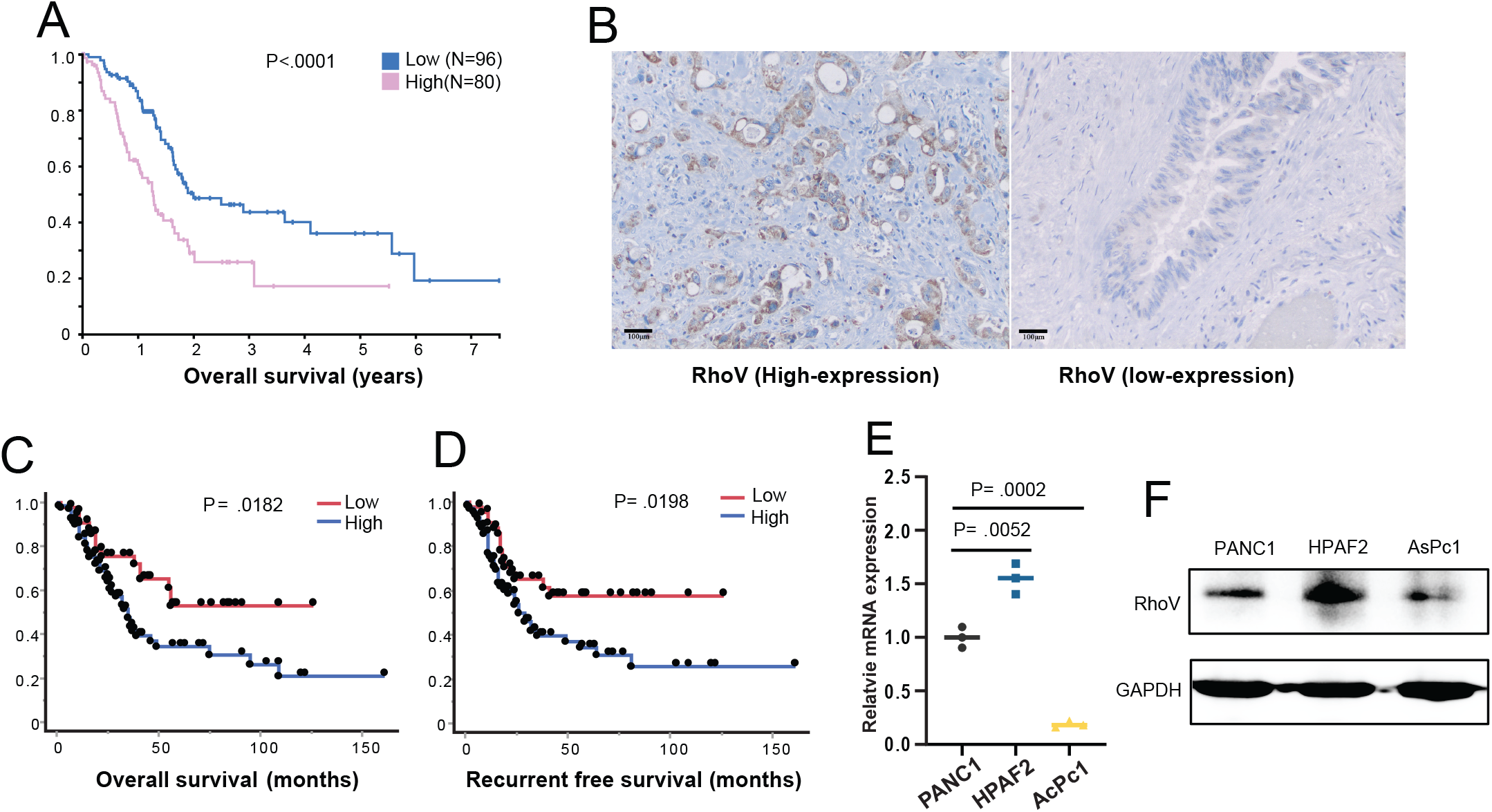
RhoV related to PDAC patient prognosis and survival rate. (A) Kaplan-Meier curve plotted high and low gene expression of RhoV showed its expression was associated with worse patient outcome (proteinatlas.org, accessed in April 2024). (B) Representative immunostain of RhoV in PDACs of high and low expression levels. (C, D) Significant difference was observed in overall survival (P=0.0182) and recurrence-free survival analysis (P=0.0198) of RhoV expression in 114 PDAC patients. (E, F) RT-qPCR and Western blotting checked both mRNA and protein levels in PANC1, HPAF2, and AsPc1 cell lines.

### Deletion of RhoV inhibits PDAC proliferation in vitro

To explore the role of RhoV in PDAC, we first knocked out the RhoV gene in PANC1 and HPAF2 cell lines with CRISPR-Cas9. We screened for stable knockout clones (T2 and T3 for both cell lines) that were confirmed at the mRNA and protein levels (Figure 2A and 2B). Then, we used MTT to investigate the effect of RhoV on PDAC cell viability. Deletion of RhoV significantly inhibited the growth of PANC1 and HPAF2 clones (Figure 2C and 2D). Concurrently performed colony formation assay also showed that the deletion of RhoV reduced the reproductive viability of PDAC cell clones (Figure 2E and 2F). In the tumor spheroid formation assay, RhoV-knockout PDAC cells (T2 and T3 clones of HPAF2 and PANC1 cells) exhibit a smaller relative diameter compared to the control, which further indicates that deletion of RhoV inhibits PDAC proliferation (Figure 2G and 2H).

**Figure 2.**
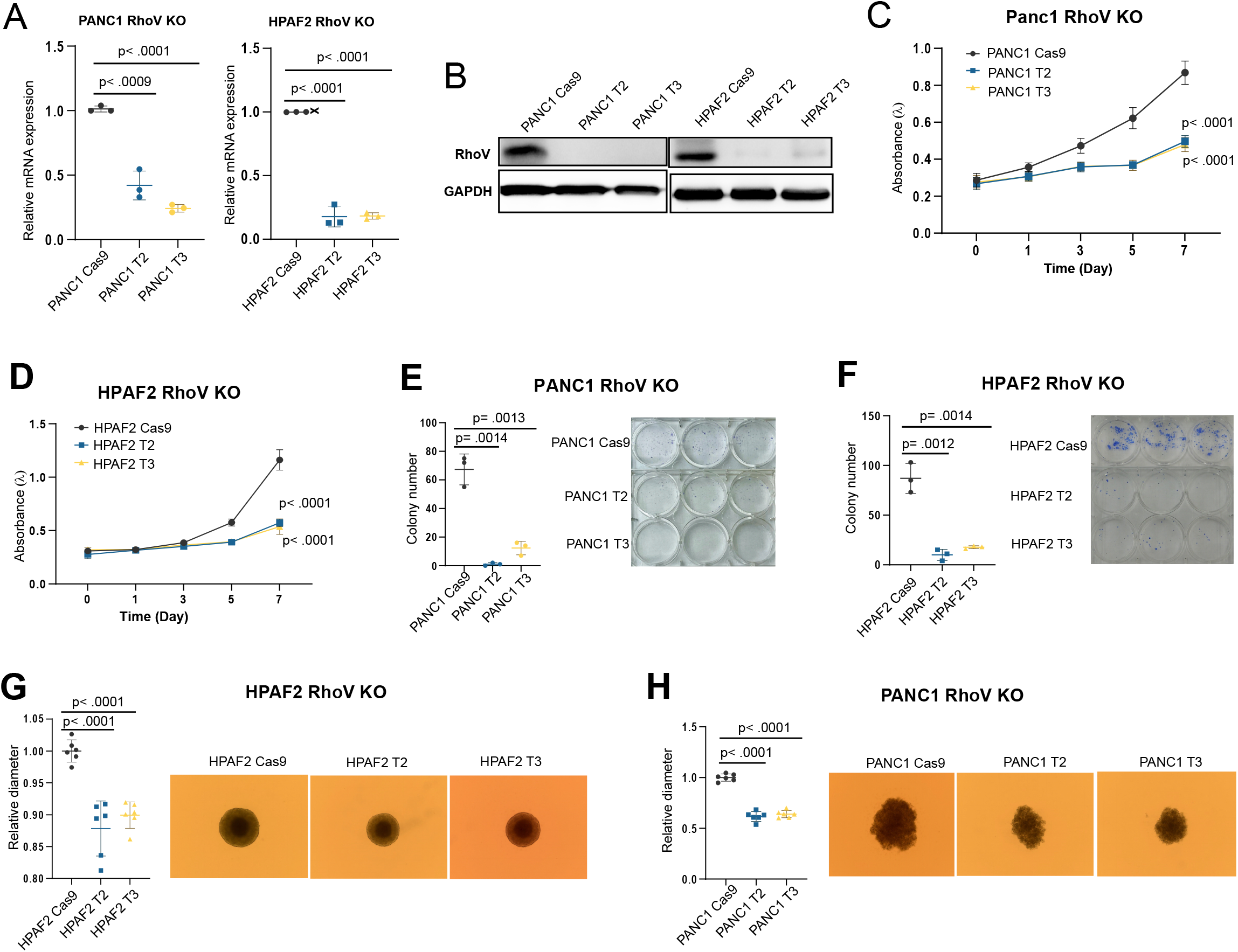
Deletion of RhoV inhibits cell proliferation in PDAC. (A) RT-qPCR examined the mRNA level in PANC1 and HPAF2 cells with stable CRISPR-Cas9 RhoV knockout or the control. (B) Western blotting results of RhoV and GAPDH showed the protein level in PANC1 and HPAF2 cells. (C, D) MTT assay showed the absorbance at 0, 1, 3, 5, 7 days in the RhoV knockout PANC1 and HPAF2 cell lines. (E, F) Colony formation was performed by Giemsa staining for 14 days. (G, H) The tumor spheroids were observed under the 4× magnification microscope on the 7th day to measure the diameter.

### Overexpressed RhoV promoted PDAC proliferation and was involved in the MAPK pathway

To further investigate the effect of RhoV on PDAC cells, we overexpressed RhoV in PANC1 and AsPc1 cell lines, which were validated at both the mRNA and protein levels (Figure 3A and 3B). In the subsequent MTT assay, PDAC cells with RhoV overexpression increased the growth capability compared to those PDAC cells with empty vector (EV) control (Figure 3C and 3D). In the colony formation assay, overexpression of RhoV promoted the growth of PDAC cells (Figure 3E and 3F). In addition, RhoV-overexpressing PDAC cells showed greater relative diameter compared to the control in the tumor spheroid assay (Figure 3G and 3H). These data suggest that RhoV promotes the proliferation of PDAC cells. Recent studies have shown that RhoV initiates the downstream JNK signaling pathway, facilitating the metastasis of triple-negative breast cancer [19]. In addition, it has been shown that in lung adenocarcinoma, RhoV contributes to tumor growth and metastasis by affecting the phosphorylation of JNK and ERK [20, 21]. To confirm whether RhoV is involved in the MAPK pathway in pancreatic adenocarcinoma, we examined the expression of the main players in the pathway, ERK, JNK, and p38. The Western blot results showed that overexpression of RhoV promoted phosphorylation of ERK, JNK, and p38 in both PANC1 and AsPc1 cell lines (Figure 3B). Taken together, these data suggested that RhoV promotes the PDAC cell growth by activating the MAPK pathway.

**Figure 3.**
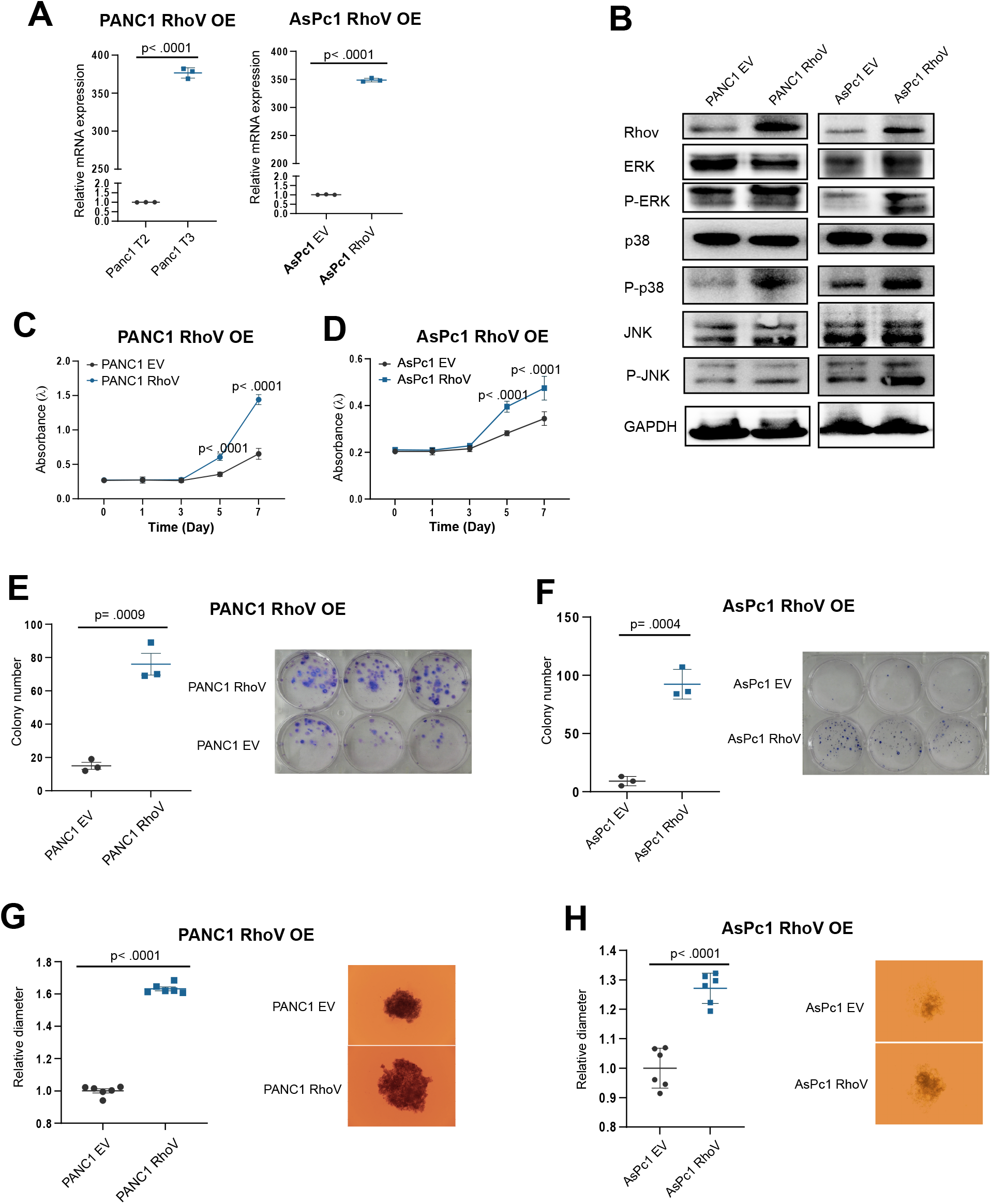
RhoV overexpression promotes cell proliferation and is involved in the MPKA pathway. (A) RT-qPCR examined the mRNA level in PANC1 and AsPc1 cells. (B) RhoV, ERK, pERK, p38, pp38, JNK, pJNK, GAPDH were determined in PANC1 and Aspc1 cells with RhoV overexpression or control. (C, D) MTT assay showed the absorbance at 0, 1, 3, 5, 7 days in the RhoV overexpression PANC1 and HPAF2 cell lines or the control. (E, F) Colony formation was performed by Giemsa staining for 14 days. (G, H) The tumor spheroids were observed under the 4× magnification microscope on the 7th day to measure the diameter.

### RhoV promotes PDAC cell migration and invasion

The Rho GTPases are known for their vital role in cell migration [23]. In the epithelial-mesenchymal transition (EMT), Rho GTPase members have been shown to be involved in the metastasis and invasion of various tumor cells [24]. To investigate whether RhoV affects the migration phenotype of PDAC cells, we performed the scratch assay and the transwell assay. In the scratch assay, the deletion of RhoV decreased the migration phenotype in PANC1 cells (Clone T2) (Figure 4A), while overexpression of RhoV promoted the migration of PDAC1 and AsPc1 cells (Figure 4B and 4C). In the transwell assay, we found a similar situation: RhoV knockout inhibited the invasion capacity of PANC1 and HPAF2 cells (Clone T2s) (Figure 4D and 4E). Subsequently, we investigated the role of RhoV in the EMT. The immunoblotting showed that knockout of RhoV increased expression of E-cadherin, but inhibited N-cadherin and Vimentin in HPAF2 cells (Clone T2) (Figure 4F). Moreover, overexpression of RhoV downregulated the expression of E-cadherin, but N-cadherin and Vimentin were increased in AsPc1 cells (Figure 4G). Our data indicated that RhoV promotes PDAC cell migration and facilitates the EMT process.

**Figure 4.**
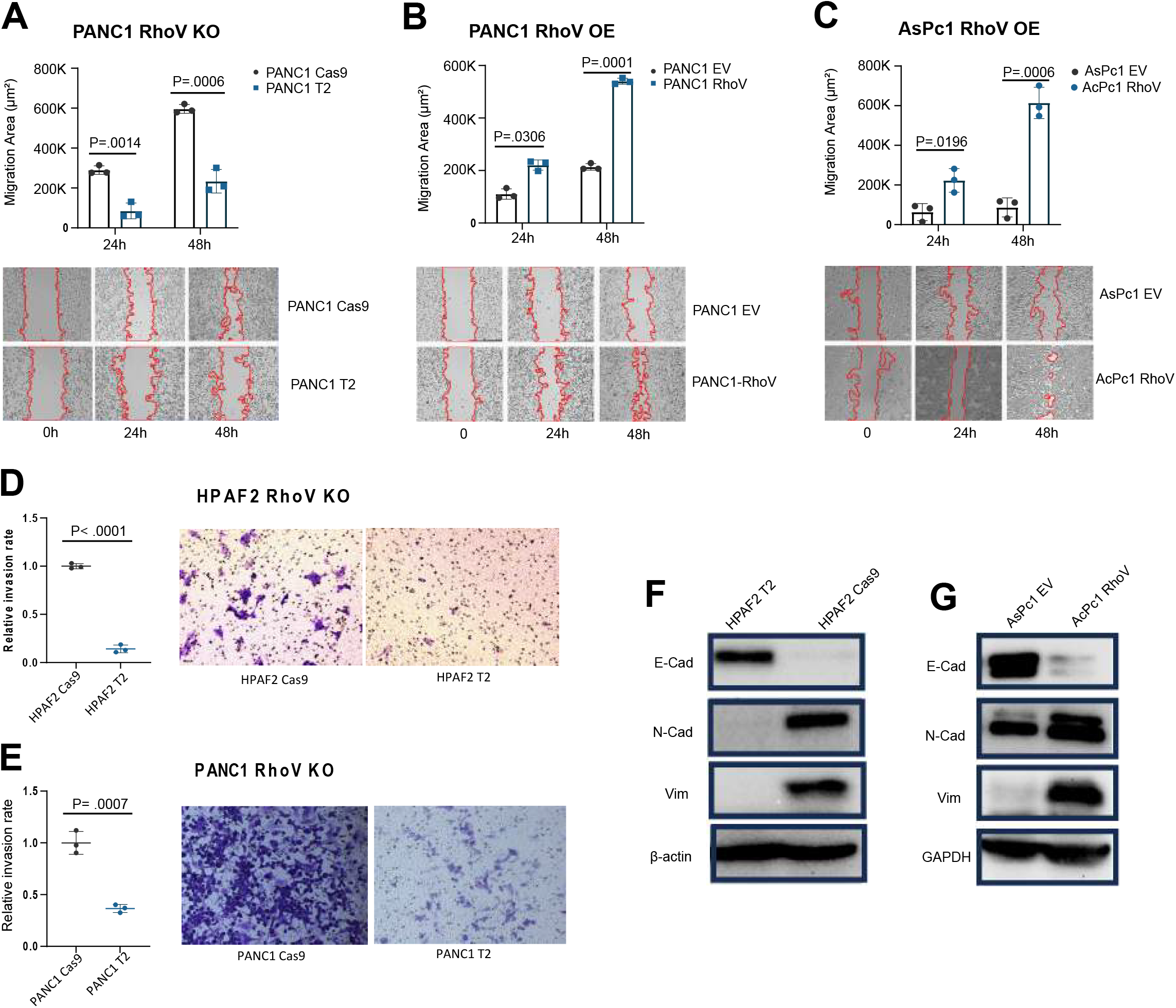
RhoV affects the migration ability of PDAC cells. (A, B) Scratch assays for PANC1 cell line with RhoV overexpression or RhoV stably knockout and the control. (C) Scratch assay for AsPc1 cell line with or without RhoV overexpression. (D, E) Transwell assay of Panc1 and HPAF2 cell lines with or without RhoV knockout. (F) E-cadherin, N-cadherin, Vimentin, and β-actin were determined in HPAF2 cells with or without RhoV knockout. (G) E-cadherin, N-cadherin, Vimentin, and GAPDH were checked in AsPc1 cells with or without RhoV overexpression.

### Modulation of RhoV affects PDAC cell growth *in vivo*

Our *in vitro* experiments have demonstrated the role of RhoV in PDAC cell proliferation, migration, and invasion. We next explored the effect of RhoV on PDAC cells *in vivo*. We made the cell line xenograft (CDX) mouse model by subcutaneously injecting the stable RhoV-modulated PDAC cells in the lower right flank (Figure 5A). Similar to what we observed in the *in vitro* experiments, overexpression of RhoV also promoted the AsPc1 cell growth *in vivo* (Figure 5B) compared to the empty vector control. Overexpression of RhoV in AsPc1 cells increased the mouse tumor burden as quantified by the ratio of tumor size to the mouse body weight (Figure 5C). Conversely, knockout of RhoV (Clone T2) inhibited tumor growth and reduced the tumor burden significantly compared to the Cas9 control (Figure 5D and 5E). From the perspective of histology, the tumors from PDAC cells with RhoV knockout (Clone T2) showed less cell density and less nuclear pleomorphism than those of the control group (Figure 5F). Our findings confirmed what was observed *in vitro* that RhoV promotes PDAC progression *in vivo*.

**Figure 5.**
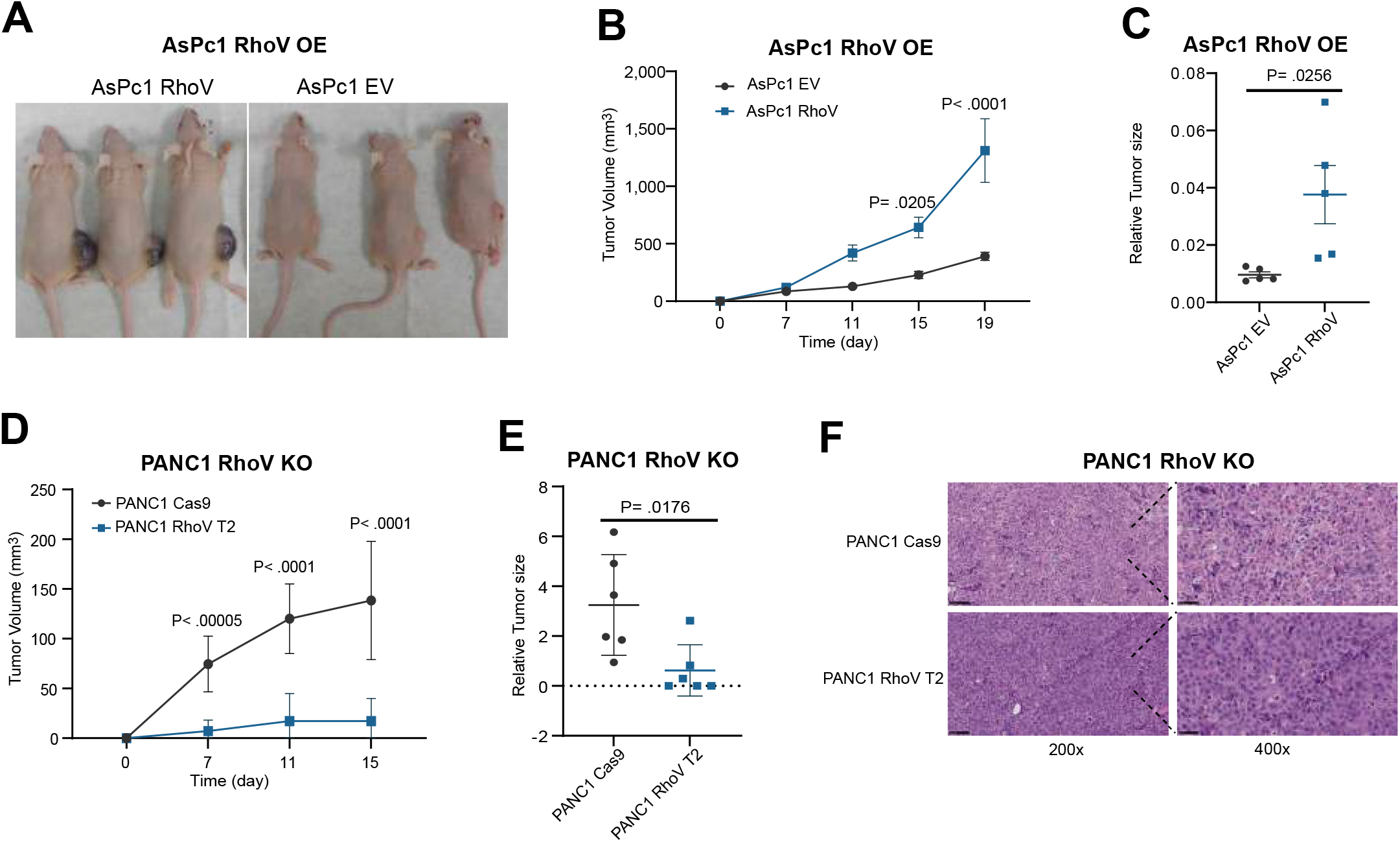
RhoV promotes PDAC growth in vivo. (A) RhoV modulated PDAC cells were subcutaneously injected into the right lower flank of NOD/SCID mice, 1X10per mouse. (B, D) Tumor volume was measured from the 7th day and every 4 days, calculated by the formulation, Tumor Volume = (Length x Width x Width)/2. (C) The relative tumor size was calculated by tumor weight over the body weight for 19 days: n = 5 in AsPc1 with or without RhoV overexpression. (E) The relative tumor size was calculated by tumor volume over the body weight for 15 days; n = 5 in PANC1 RhoV knockout; n = 6 in PANC1 Cas9. (F) Representative hematoxylin and eosin staining images (200× and 400×) in PANC1 RhoV knockout and the control group.

## Discussion

In this study, we comprehensively investigated the role of RhoV in pancreatic ductal adenocarcinoma (PDAC) using a combination of surgically resected PDAC samples, *in vitro* cellular models, and *in vivo* xenograft assays. We demonstrated that high RhoV expression in PDACs is significantly associated with worse overall survival and recurrence-free survival in patients. Functional studies revealed that RhoV promotes PDAC cell proliferation, colony and spheroid formation, and enhances migratory and invasive capacities. Mechanistically, RhoV overexpression activated key components of the MAPK signaling pathway, demonstrated by increased phosphorylation of ERK, JNK, and p38. In addition, RhoV overexpression promoted the process of EMT, while RhoV-knockout reversed mesenchymal phenotypes. These findings collectively establish RhoV as a putative oncogenic driver in PDAC progression.

Although Rho GTPases are broadly recognized for their roles in cytoskeletal dynamics and cell motility [23,24], emerging evidence suggests their functions also extend to transcriptional regulation, survival signaling, and metabolic adaptation [13]. The present data support the notion that RhoV functions as a central mediator linking extracellular cues to intracellular MAPK pathway activation in PDAC. This is consistent with prior studies in breast and lung cancers, where RhoV facilitated tumor progression through JNK and ERK activation [19–21]. The observation that RhoV modulated EMT markers such as E-cadherin, N-cadherin, and Vimentin further implicates RhoV as a regulator of epithelial plasticity, which is known to underpin metastatic competence [24]. It is plausible that RhoV contributes to PDAC aggressiveness not only by promoting proliferation but also by facilitating tumor cell dissemination through EMT induction and enhanced matrix invasion. However, the precise upstream mechanisms regulating RhoV expression in PDAC remain undefined and warrant further investigation.

Molecular subtyping has highlighted the heterogeneity of PDAC, with the basal-like subtype characterized by particularly poor prognosis, high proliferative index, and resistance to chemotherapy [22]. Notably, analyses of publicly available datasets and immunohistochemical profiling revealed that RhoV is preferentially enriched in basal-like PDAC tumors [17]. This raises the possibility that RhoV may contribute to the aggressive phenotype of this molecular subtype. Given the limited therapeutic options for basal-like PDAC, RhoV could represent a candidate biomarker for stratification and a potential target for intervention. Future studies should explore whether targeting RhoV selectively affects basal-like tumors or whether it holds broader therapeutic relevance across PDAC subtypes.

Despite advances in targeting KRAS mutations, including the development of selective KRAS G12C inhibitors [10], the clinical benefit in PDAC remains modest [25], partly due to the emergence of adaptive resistance mechanisms [11]. The identification of RhoV as a facilitator of MAPK pathway activation suggests that RhoV may act as a bypass signaling hub enabling tumor cells to sustain proliferation despite upstream KRAS blockade. This aligns with the concept of pathway redundancy and network rewiring observed in KRAS inhibitor resistance [12]. Recent phosphoproteomic and transcriptomic studies have delineated how KRAS and ERK signaling orchestrate complex adaptive programs that sustain tumor growth even in the presence of targeted inhibitors [26, 27]. Accordingly, combinatorial targeting of KRAS and RhoV—or the downstream MAPK effectors activated by RhoV—may represent a rational therapeutic strategy to overcome resistance and improve treatment durability. Further preclinical studies are warranted to test this hypothesis in relevant KRAS-driven PDAC models.

